# Acetyltransferases GCN5 and PCAF are required for B lymphocyte maturation in mice

**DOI:** 10.1101/2021.12.11.472222

**Authors:** Valentyn Oksenych, Dan Su, Jeremy A. Daniel

**Affiliations:** The NNF Center for Protein Research, Faculty of Health and Medical Sciences, University of Copenhagen, Blegdamsvej 3B, 2200 Copenhagen, Denmark; Department for Cancer Research and Molecular Medicine (IKOM), Norwegian University of Science and Technology, Laboratory Center, Erling Skjalgssons gate 1, 7491 Trondheim, Norway; KG Jebsen Centre for B Cell Malignancies, Institute of Clinical Medicine, University of Oslo, N-0316 Oslo, Norway; Institute of Clinical Medicine, University of Oslo, 0318 Oslo, Norway

**Keywords:** KAT2A, KAT2B, mice, acetyltransferase, B cell, lymphocyte, class switching

## Abstract

B lymphocyte development has two DNA recombination processes: V(D)J recombination of the immunoglobulin (*Igh*) gene variable region and class switching of the *Igh* constant regions from IgM to IgG, IgA, or IgE. V(D)J recombination is required for successful maturation of B cells from pro-B to pre-B to immature-B and then to mature B cells in the bone marrow. CSR occurs outside of the bone marrow when mature B cells migrate to peripheral lymphoid organs, such as spleen and lymph nodes. Both V(D)J recombination and CSR depend on an open chromatin state that makes DNA accessible to specific enzymes, recombination activating gene (RAG), and activation-induced cytidine deaminase (AID). Acetyltransferases GCN5 and PCAF possess redundant functions acetylating histone H3 lysine 9 (H3K9). Here, we generated a mouse model that lacks both GCN5 and PCAF in B cells. We found that double-deficient mice possess low levels of mature B cells in the bone marrow and peripheral organs, an accumulation of pro-B cells in bone marrow, and reduced CSR levels. We conclude that both GCN5 and PCAF are required for B cell development *in vivo*.

## 1. Introduction

Development of B lymphocytes starts in the bone marrow where progenitor (pro)-B cells using recombination-activating genes (RAG) generate DNA double-strand breaks (DSBs) and initiate the V(D)J recombination (1). In maturating B cells, the V(D)J recombination process is genetic recombination of *variable (V), diversity (D)*, and *joining (J)* gene segments arranging into a newly formed VDJ part of immunoglobulin gene (*Ig*) (2–6). Following the V(D)J recombination, B cells develop from pro-B cells expressing specific markers cluster of differentiation 19 (CD19), B220/CD45, and CD43 (CD19+B220+CD43+) to pre-B cells (CD19+B220+CD43-), immature B (CD19+B220+IgM+, low immunoglobulin M, IgM) and mature B (CD19+B220+IgM+, high IgM) cells in bone marrow (2). Mature B lymphocytes leave the bone marrow and through the blood migrate to periphery populating spleen and lymph nodes.

Then, mature B cells initiate another DNA recombination process to change the constant regions of immunoglobulin genes referred to as class switch recombination (CSR). During the CSR in mice, IgM is replaced by IgG3, IgG1, IgG2a, IgG2b, IgE, or IgA (2). The CSR is initiated by non-productive transcription known as a germ-line transcription (GLT) which is needed to separate two DNA strands. Single-stranded DNA is then targeted by activation-induced cytidine deaminase (AID), a B lymphocyte-specific enzyme deaminating cytosine to uracil (C to U). Then uracil DNA N-glycosylase (UNG) removes the uracil from DNA leading to the single-strand break formation (SSB)(7). Two SSBs facing each other then form a DSB and allow recombination (2–5, 8). Both V(D)J and CSR can be regarded as processes following fundamentally similar strategies of genomic recombination (9, 10).

The DSBs formed during the V(D)J recombination and CSR are recognized, processed, and repaired by the non-homologous end-joining pathway (NHEJ) and initiate more complex signaling and chromatin modification pathway known as DNA damage response (DDR) (2–5). The NHEJ is initiated when Ku70 and Ku80 heterodimer (Ku) recognizes and binds the DSBs. Ku serves as a platform for downstream factors including DNA-dependent protein kinase catalytic subunit (DNA-PKcs), X-ray repair cross-complementing protein 4 (XRCC4), XRCC4-like factor (XLF), Paralogue of XRCC4 and XLF (PAXX), a modulator of retrovirus infection (MRI), and DNA ligase 4 (LIG4) (3–5, 11). There are additional factors that are sometimes optional for NHEJ, including Artemis with nuclease activity required for processing hairpin-sealed DNA ends and overhangs (2, 5, 12).

One type of DDR pathway acts downstream of ataxia telangiectasia mutated (ATM) protein kinase, which is activated by DSBs and then phosphorylates multiple substrates, including NHEJ and DDR factors. ATM phosphorylates histone H2AX, which in turn recruits mediator of DNA damage checkpoint 1 (MDC1) and facilitates the accumulation of really interesting new gene (RING) finger motif (RNF) 8 and RNF168 ubiquitin-ligases, and then the p53-binding protein (53BP1). Phosphorylation of H2AX is related to the acetylation of histones, including histone H3K9. In particular, histone acetylation relies on the ATM-dependent H2AX phosphorylation and SWI/SNF chromatin remodeling factors (13). Acetylation of histone H3K9 is mediated by GCN5 and PCAF (13, 14).

There is a complex genetic interaction associated with the functional redundancy between the NHEJ factors (3, 4), including the following pairs: DNA-PKcs/XLF (5, 15–17), PAXX/XLF (18–24), and MRI/XLF (25, 26). Moreover, there is a genetic interaction between NHEJ and DDR pathway factors, for example, ATM/XLF and H2AX/XLF (27), MDC1/XLF (28), RNF8/XLF and RNF168/XLF (29), 53BP1/XLF (30, 31), and more (3, 4). Moreover, acetyltransferases GCN5 and PCAF are redundant in promoting histone H3 lysine K9 acetylation (32).

General control non-depressible 5 (GCN5) acetyltransferase is also known as lysine acetyltransferase (KAT) 2A. Germline inactivation of GCN5 in mice resulted in early embryonic lethality due to the role of the protein in neurogenesis (32). GCN5 is functionally redundant with another acetyltransferase, KAT2B, also referred to as p300/CBP-associated factor (PCAF). While inactivation of *Pcaf* gene in mice has no detectable phenotype, double knockout of *Gcn5/Pcaf* genes resulted in even earlier embryonic lethality than in *Gcn5^-/-^* mice (32). Because histone H3K9 acetylation works downstream of ATM and H2AX in DDR (13), one could propose that GCN5, PCAF, or both enzymes are required for lymphocyte development *in vivo*. However, embryonic lethality of *Gcn5^-/-^* and *Gcn5^-/-^ Pcaf^-/-^* mice (32) challenged the studies. To overcome the obstacle, we developed a complex mouse model, when *Pcaf* gene was germline-inactivated (32), while floxed *Gcn5* gene (33) was conditionally-inactivated in B cell lineages by CRE recombinase expressed under *Cd19* promoter (34). To sort out CRE-positive and CRE-negative cells, we used *Rosa26-stop-YFP* knockin which only expressed YFP following the CRE activation (35).

Here, we found that GCN5 and PCAF acetyltransferases are functionally redundant during early B cell maturation, while GCN5 is required for robust CSR.

## 2. Materials and Methods

### 2.1. Mice

*Gcn5^f/f^* (33), *Pcaf^+/-^* (32), *Cd19^Cre+^* (34) (# 006785; The Jackson Laboratory, Bar Harbor, ME, USA), *Rosa26-stop-YFP^+^* (35)(# 006148, The Jackson Laboratory, Bar Harbor, ME, USA), and *Aid^-/-^* (36) mice were previously described. Mice used for experiments were between 8-12 weeks of age. All experiments were performed in compliance with the Danish Working Environment Authority, the Danish Animal Experiment Inspectorate, the Department of Experimental Medicine (University of Copenhagen), and the Animal Resources Care Facility of Norwegian University of Science and Technology (NTNU, Trondheim, Norway).

### 2.2. Flow cytometry

Flow cytometry experiments were performed as we described earlier (25, 28, 37–39). In particular, we used the fluorescent antibodies recognizing the proteins described below. B220 (PE-CF594, FITC, Alexa Fluor 700; all clone RA3-6B2, BD Bioscience, Franklin Lakes, NJ, USA). IgM (PerCP-eFluor 710, APC-eFluor 780, APC, FITC; all clone II/41, eBioscience, Santa Clara, CA, USA). IgG1 (PE, MOPC-21, Biolegend, SanDiego, CA, USA). IgG3 (PE, MG3-35, Biolegend, San Diego, CA, USA). CD3-APC (Biolegends, USA, #100312). CD19 (Alexa Fluor 700, APC eFluor 780, both clone 1D3, eBioscience, Santa Clara, CA, USA). CD43 (APC and PE-Cy7, both clones S7, BD Bioscience, Franklin Lakes, NJ, USA).

### 2.3. Class switch recombination

The CSR was performed as we described earlier (15, 27, 38–42).

### 2.4. Western blot

The western blot procedures were performed as we described earlier (28, 37, 38, 43). Briefly, the cells were lysed for 30 minutes on ice in Radioimmunoprecipitation assay (RIPA) buffer (Sigma Aldrich, St. Louis, MO, USA, #R0278) supplemented with cOm-plete™ EDTA-free Protease Inhibitor Cocktail (Sigma Aldrich, #11873580001). Proteins were analyzed by 4-12% Bis-Tris NuPAGE gels (Invitrogen, Carlsbad, CA, USA, #NP0322), transferred to PDVF membranes (GE Healthcare, Boston, MA, USA, #GE10600023), and probed with indicated antibodies. Rat anti-AID (1:500, Active Motif, Carlsbad, CA, USA, #39886); mouse anti-GCN5, clone A-11 (1:500, Santa Cruz Biotechnology, Dallas, TX, USA, #sc-365321); rabbit anti-PCAF, clone C14G9 (1:1000, Cell Signaling Technology, Leiden, The Netherlands, #3378); rabbit anti-histone H3 (1:1000, Abcam, Cambridge, UK, #ab1791); rabbit anti-histone H3 acetyl K9 (1:500, Abcam, Cambridge, UK, #ab32129).

### 2.5. Statistics

We performed statistical analyses with one-way ANOVA using GraphPad Prism 8.0.1.244 (San Diego, CA, USA). In the tests, p values less than 0.05 were defined as significant, i.e. *p<0.05; **p<0.01; ***p<0.001; and ****p<0.0001.

## 3. Results

### 3.1. Generation of mice lacking GCN5 and PCAF in B cells

Combined inactivation of *Gcn5* and *Pcaf* genes in mice results in embryonic lethality (32). To overcome this challenge, we designed a complex genetic model when floxed *Gcn5* gene is conditionally inactivated in B cell lineages by CRE enzyme under the *Cd19* promoter (*Cd19^Cre+^*) (34). To sort out the cells with activated CRE, we used a model with knocked-in yellow fluorescent protein gene (*YFP*) into ROSA-26 locus. The YFP is inactive until CRE removes “STOP” signal (Rosa-26-YFP+) (35). Thus, we obtained *Gcn5^f/f^Pcaf^-/-^ Cd19^+/Cre^YFP^+^* mice and simpler controls. Further in the text, we will skip *Cd19^+/Cre^* and *YFP+* for simplicity in most of the cases, and will refer to the mice based on the status of *Gcn5* and *Pcaf* genes, i.e. as *Gcn5^f/f^Pcaf^-/-^*, *Gcn5^f/f^*, *Pcaf^-/-^* and WT. When the CRE is active and describing sorted B cells, we indicate *Gcn5^-/-^*, a knockout status of the gene. Lack of GCN5 and PCAF, as well as H3K9 acetylation, was validated using western blot (e.g., Figure S1).

### 3.2. Mice lacking GCN5 and PCAF in B cells possess small spleens

We obtained mice with germline inactivation of *Pcaf* gene and conditional inactivation of *Gcn5* in B cells under the *Cd19* promoter (Figure 1). We found that germline inactivation of *Pcaf* gene alone has no detectable effect on mouse development, in line with the previous observation (32). Conditional inactivation of *Gcn5* gene in B cells had no visible effect on sizes of WT and *Pcaf*-deficient mice, which were 15 to 19 g on average (p>0.1433) (Figure 1A). However, inactivation of *Gcn5* resulted in smaller spleens in mice (*Gcn5^-/-^*, 54 mg), when compared to WT (69 mg) and *Pcaf^-/-^* (72 mg) mice. Combined inactivation of *Pcaf* and *Gcn5* in B cells resulted in even smaller spleens (*Gcn5^-/-^Pcaf^-/-^*, 29mg, p<0.0001). Spleens of mice without CRE activity with *Gcn5* gene being floxed and functional (*Gcn5^f/f^Pcaf^-/-^*, 67 mg), were comparable in size to the ones of WT (Figure 1 B, C).

**Figure 1.**
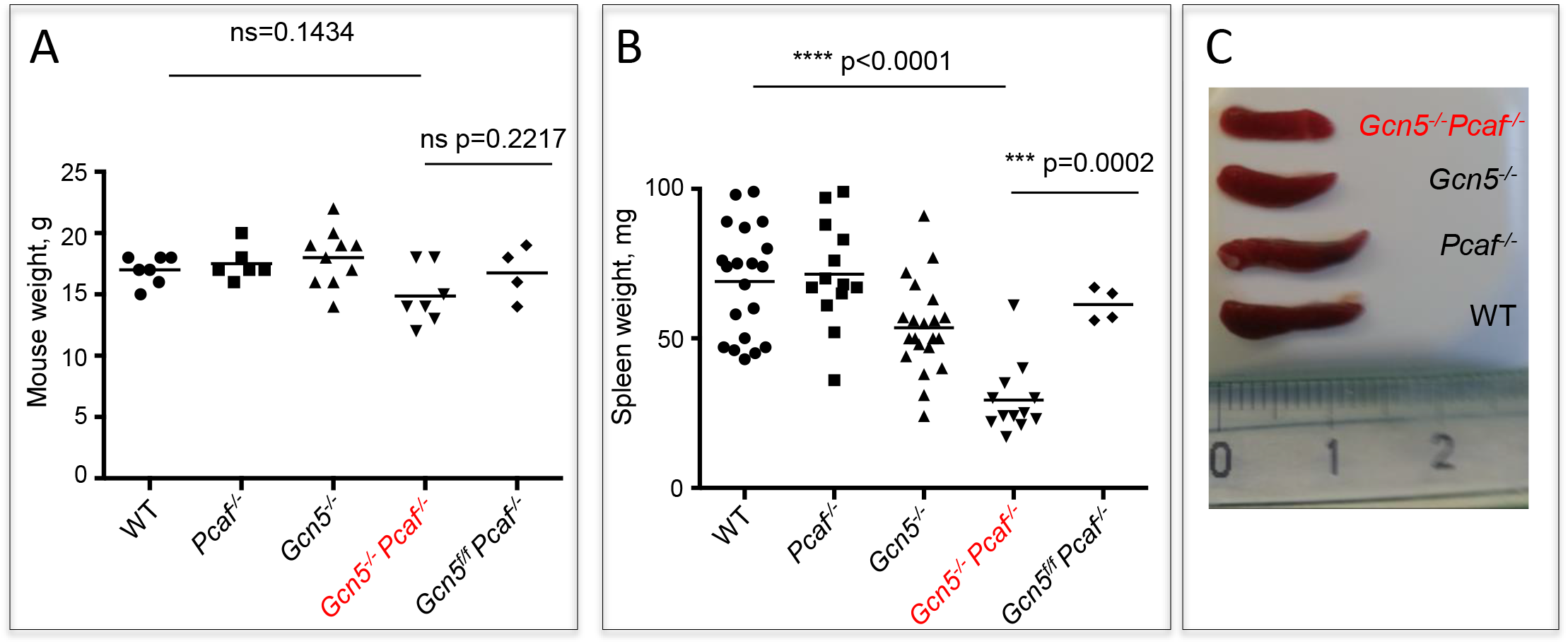
Generation of mice with germline inactivation of *Pcaf* and conditional inactivation of *Gcn5* in B cell lineages. (A) Sizes of 8 weeks-old mice of indicated genotype are similar (p>0.1433). (B) Size of spleens from mice of indicated genotypes. WT vs *Pcaf^-/-^*, n.s. p=0.9559; WT vs *Gcn5^-/-^*, *p=0.0192; WT vs *Gcn5^-/-^Pcaf^-/-^*, ****p<0.0001; *Pcaf^-/-^* vs *Gcn5^-/-^*, *p=0.0119; *Pcaf^-/-^*vs *Gcn5^-/-^Pcaf^-/-^*, ****p<0.0001; *Gcn5^-/-^* vs *Gcn5^-/-^Pcaf^-/-^*, ***p=0.0008. (C) Example of spleens of indicated genotypes. *Gcn5^-/-^* indicates *Cd19^Cre^*-dependent inactivation of *Gcn5* in B cell lineages.

### 3.3. Mice lacking GCN5 and PCAF in B cells possess delayed B lymphocyte development

To detect mature *Gcn5^-/-^Pcaf^-/-^* B cells, we identified B220+IgM+ cells in the spleen using flow cytometry (Figure 2 A, B). Inactivation of *Pcaf* gene alone did not affect B cell proportions in the spleen (58%) when compared to WT mice (52%, p=0.4777) (Figure 2A). Inactivation of *Gcn5* alone resulted in an insignificant reduction of mature splenocytes when compared to WT (*Gcn5^-/-^*, 46%, p<0.2532), although *Gcn5^-/-^Pcaf^-/-^* mice were found to have a significantly less B cell frequency in the spleen (21%, p<0.0001). Similarly, the numbers of *Gcn5^-/-^Pcaf^-/-^* B splenocytes was the lowest (3,4 million), while the number of *Gcn5^-/-^* B cells (27 million) was also reduced when compared to *Pcaf^-/-^* (54 million, **p=0.0065) and WT (44 million, *p=0.0214) controls (Figure 2B).

**Figure 2.**
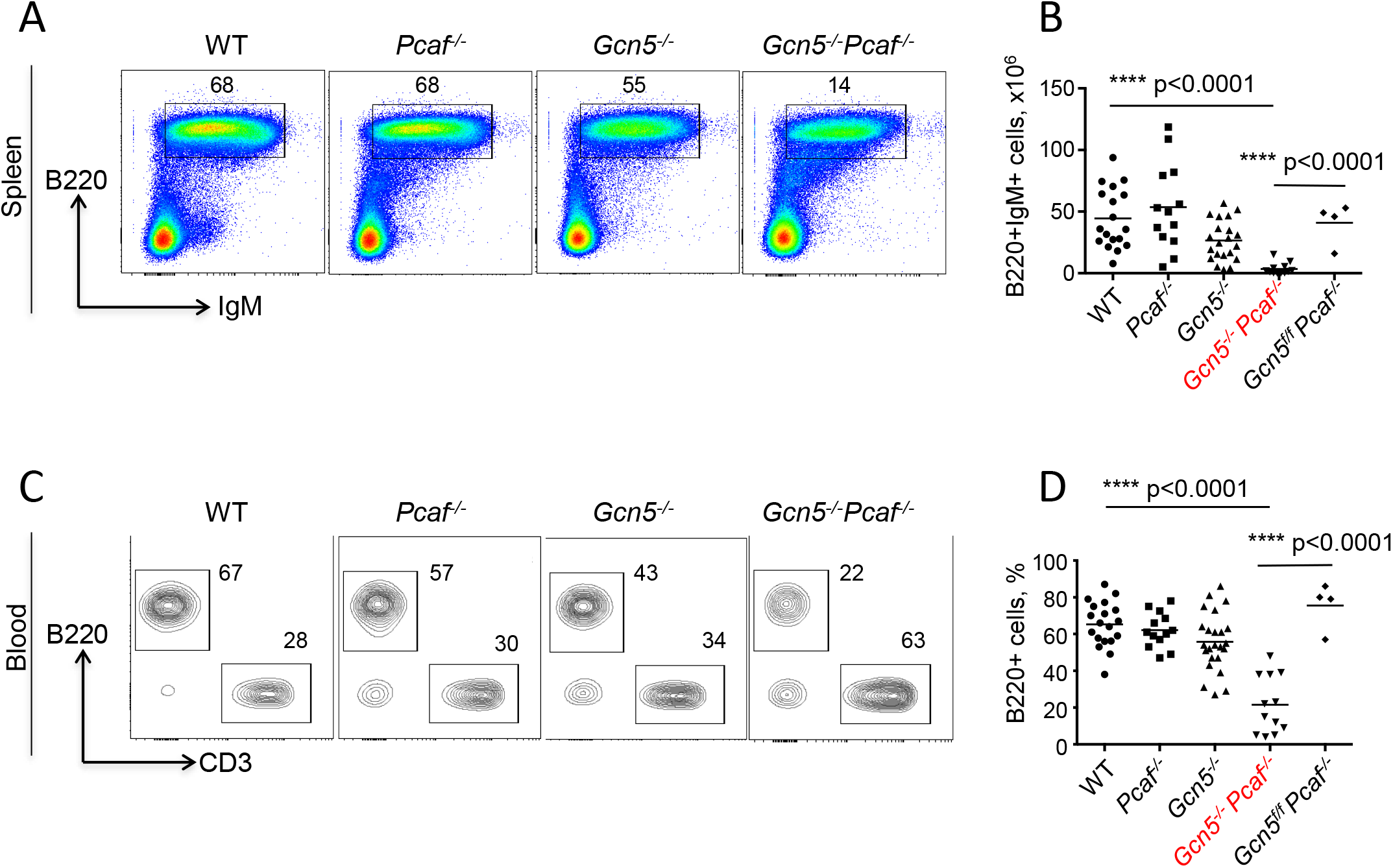
Reduced levels of mature B lymphocytes in spleens and blood of 8-12-weeks-old mice of indicated genotypes. (A) Proportions of B220+IgM+ mature B cells in spleen. WT vs *Pcaf^-/-^*, n.s. p=0.4777; WT vs *Gcn5^-/-^*, ns, p=0.2532; WT vs *Gcn5^-/-^Pcaf^-/-^*, ****p<0.0001; *Pcaf^-/-^* vs *Gcn5^-/-^*, *p=0.0119; *Pcaf^-/-^* vs *Gcn5^-/-^Pcaf^-/-^*, ****p<0.0001; *Gcn5^-/-^* vs *Gcn5^-/-^Pcaf^-/-^*, ***p<0.0001. (B) Summary of experiments as shown in (A), number of cells. WT vs *Pcaf^-/-^*, n.s., p=0.9212; WT vs *Gcn5^-/-^*, *p=0.0214; WT vs *Gcn5^-/-^Pcaf^-/-^*, ****p<0.0001; *Pcaf^-/-^* vs *Gcn5^-/-^*, **p=0.0053; *Pcaf^-/-^* vs *Gcn5^-/-^Pcaf^-/-^*, ****p<0.0001; *Gcn5^-/-^* vs *Gcn5^-/-^Pcaf^-/-^*, *p<0.0269. (C) Proportions of B220+ B cells and CD3+ T cells in blood. (D) Summary of several experiments from (C) reflecting proportions of B cells in blood. WT vs *Pcaf^-/-^*, n.s. p=0.9196; WT vs *Gcn5^-/-^*, n.s., p=0.1204; WT vs *Gcn5^-/-^Pcaf^-/-^*, ****p<0.0001; *Pcaf^-/-^* vs *Gcn5^-/-^*, n.s., p=0.5398; *Pcaf^-/-^*vs *Gcn5^-/-^Pcaf^-/-^*, ****p<0.0001; *Gcn5^-/-^* vs *Gcn5^-/-^Pcaf^-/-^*, ***p<0.0001. *Gcn5^-/-^* and *Gcn5^-/-^Pcaf^-/-^* indicate *Cd19^Cre^*-dependent inactivation of *Gcn5* in B cell lineages.

### 3.4. Inactivation of Gcn5 and Pcaf results in a reduced proportion of B cells in the blood

To detect mature B cells in the blood, we used B220 markers (Figure 2C, D). Inactivation of *Pcaf* alone resulted in 62% of B cells after red blood cells were lysed, which was comparable to WT mice with 65% of B cells in the blood (p=0.92). Inactivation of *Gcn5* gene alone resulted in a modest reduction of B cell proportion to 56% (p=0.12), while combined inactivation of *Gcn5* and *Pcaf* led to even lower B cell levels in blood (22%, p<0.0001). Levels of B cells in blood of control mice without CRE recombinase expression, when *Gcn5* gene was functional (*Gcn5^f/f^Pcaf^-/-^*) were comparable to WT mice (78%). We conclude that GCN5 and PCAF are both required and functionally redundant for B cell development in mice.

One reason for low *Gcn5^-/-^Pcaf^-/-^* B cell count in spleen and blood in mice could be cell death following normal development of B cells in bone marrow and migration to periphery. Another option could be blocked or delayed maturation of B cells in bone marrow during the earlier developmental stages. To test the latter possibility, we analyzed B cells in bone marrow of the mice (Figure 3).

**Figure 3.**
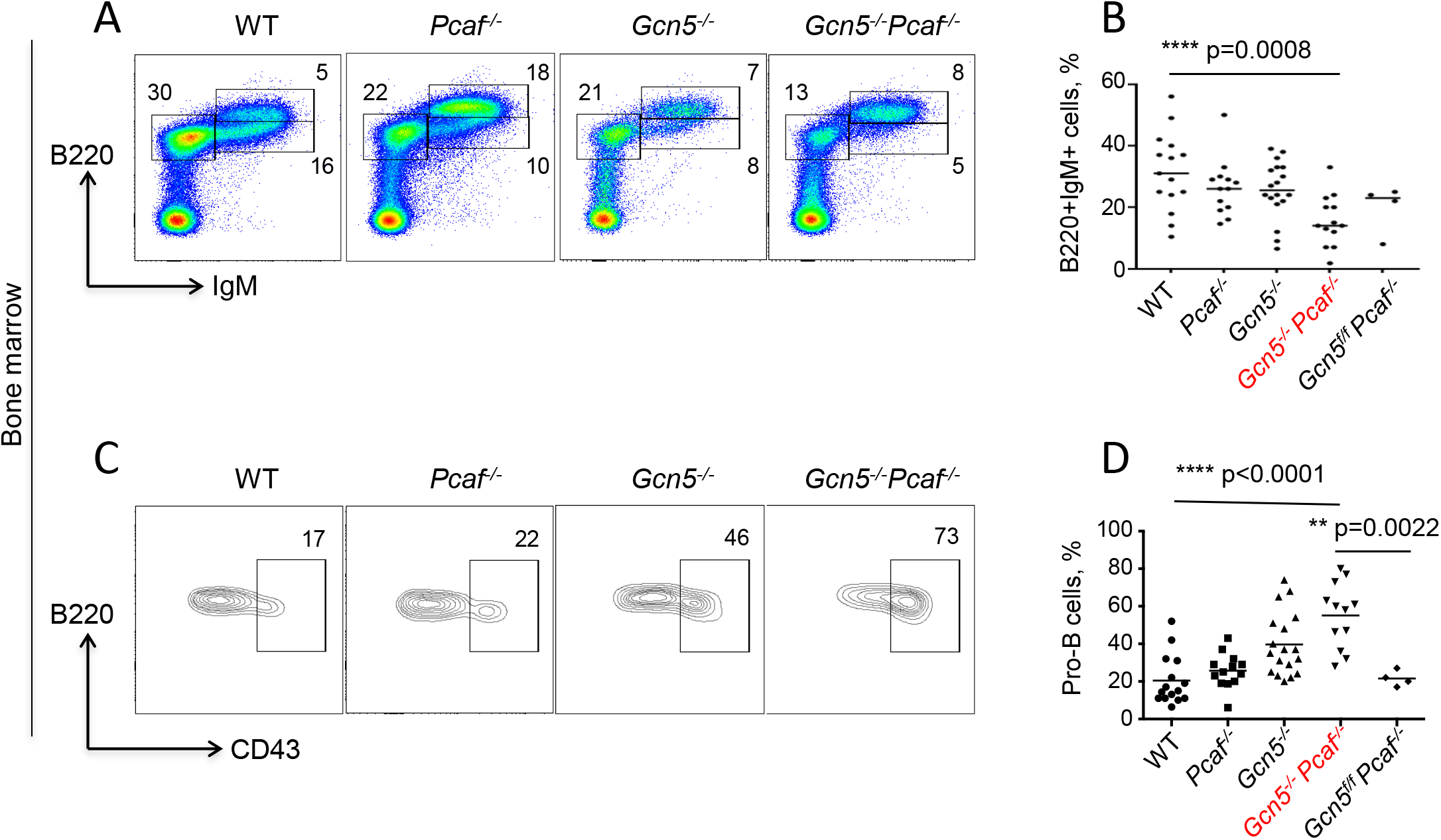
Delayed development of GCN5/PCAF-deficient B cells in bone marrow. Flow cytometry analyses of developing B lymphocytes in bone marrow of 8-12-week-old mice of indicated genotypes. (A) Examples of B220+IgM- (pro-B and pre-B), B220+IgM+low (immature B) and B220+IgM+high (mature B) cell populations. (B) Summary of several experiments from (A) indicating B220+IgM+ cells. WT vs *Pcaf^-/-^*, n.s. p=0.6445; WT vs *Gcn5^-/-^*, n.s., p=0.4432; WT vs *Gcn5^-/-^Pcaf^-/-^*, ***p=0.0008; *Pcaf^-/-^* vs *Gcn5^-/-^*, n.s., p=0.9997; *Pcaf^-/-^* vs *Gcn5^-/-^Pcaf^-/-^*, *p<0.0138; *Gcn5^-/-^* vs *Gcn5^-/-^Pcaf^-/-^*, *p<0.0211 (C) CD43+ (pro-B cells) and CD43- (pre-B cells) gated from B220+IgM-populations in (A). (D) Summary of several experiments detecting B220+IgM-CD43+ (pro-B) cells.. WT vs *Pcaf^-/-^*, n.s. p=0.7825; WT vs *Gcn5^-/-^*, **p=0.0022; WT vs *Gcn5^-/-^Pcaf^-/-^*, ****p<0.0001; *Pcaf^-/-^* vs *Gcn5^-/-^*, n.s., p=0.0505; *Pcaf^-/-^* vs *Gcn5^-/-^Pcaf^-/-^*, ****p<0.0001; *Gcn5^-/-^* vs *Gcn5^-/-^Pcaf^-/-^*, *p<0.0322 *Gcn5^-/-^* and *Gcn5^-/-^Pcaf^-/-^* indicate *Cd19^Cre^*-dependent inactivation of *Gcn5* in B cell lineages.

### 3.5. Inactivation of Gcn5 and Pcaf results in accumulation of pro-B cells in bone marrow

To characterize B cell maturation in bone marrow, we followed the expression of B220 (B220+IgM-) and IgM (B220+IgM+) on the lymphocyte surface. Inactivation of *Gcn5* or *Pcaf* resulted in an insignificant decline of B220+IgM+ population (26%, 5-6 million) when compared to WT (32%, 8 million, p=0.44)(Figure 3 A, B). Combined inactivation of *Gcn5* and *Pcaf* resulted in an additional reduction of mature B cells in bone marrow (3 million, 16%) (Figure 3 A, B).

We further focused on B220+IgM-populations by determining the CD43+ (pro-B cells) and CD43- (pre-B cells). Proportion of early-stage pro-B cells increased from WT mice (20% on average) and *Pcaf^-/-^* mice (26%, p=0.78) to *Gcn5^-/-^* (39%, **p=0.0022) to *Gcn5^-/-^Pcaf^-/-^* (55%, p<0.0001) (Figure 3 C, D). However, the number of pro-B cells in bone marrow estimated using our method of cell extraction was rather stable, with 0.8 million for WT (n=13) and about 1.3 million for *Gcn5^-/-^* (n=13), *Pcaf^-/-^* (n=13), and *Gcn5^-/-^Pcaf^-/-^* (n=5) cells (n.s., p>0.3830). It suggested that the proportion of *Gcn5^-/-^Pcaf^-/-^* pro-B cells was increased because the total number of mature B cells was reduced (Supplemental Figure S3 and Figure 3B).

We conclude that GCN5 and PCAF are required for the maturation of B cells from the pro-B cell stage to pre-B and later to mature B cells.

### 3.6. GCN5 is required for robust class switch recombination

The CSR relies on the ATM-dependent DDR (2, 4, 5, 27). Because H3K9 acetylation works downstream of H2AX phosphorylation, and GCN5/PCAF might work downstream of ATM/ATR/DNA-PKcs, we tested if the CSR depends on GCN5 and PCAF (Figure 4). We purified B splenocytes from 8 to 12 weeks old mice and stimulated the CSR from IgM to IgG3 using established protocols (40, 42). We focused on matched pairs of *Gcn5^f/f^* (the functional equivalent of WT cells) and *Gcn5^-/-^*, as well as *Gcn5^f/f^Pcaf^-/-^* (the functional equivalent of *Pcaf^-/-^*) and *Gcn5^-/-^Pcaf^-/-^* cells. We used *Aid^-/-^* cells as a CSR-deficient control to detect an experimental background (Figure 4). Inactivation of *Pcaf* alone had no effect on CSR levels (WT vs *Gcn5^f/f^Pcaf^-/-^*, p>0.96). Contrary, the inactivation of *Gcn5* resulted in a reduction of CSR from about 14% in WT and *Gcn5^f/f^* cells to 6% in *Gcn5^-/-^* cells, *p<0.0008 (Figure 4). Combined deletion of *Pcaf* and *Gcn5* resulted in a similar reduction from 12% in *Gcn5^f/f^Pcaf^-/-^* cells to 6% in *Gcn5^-/-^Pcaf^-/-^* cells, **p=0.0080. We conclude that GCN5 is required for robust CSR to IgG3 because additional inactivation of *Pcaf* did not affect CSR levels when compared *Gcn5^-/-^* and *Gcn5^-/-^Pcaf^-/-^* B cells, p>0.9999 (Figure 4).

**Figure 4.**
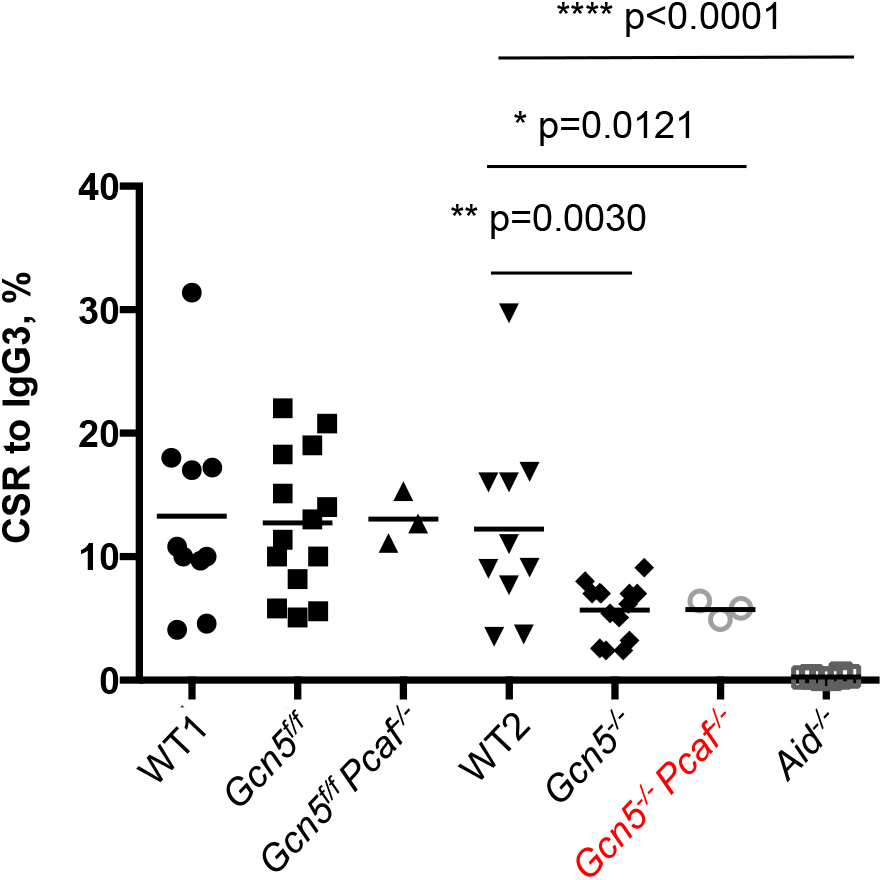
Class switch recombination of stimulated mature B cells from IgM to IgG3. *Gcn5^-/-^* and *Gcn5^-/-^Pcaf^-/-^* indicate *Cd19^Cre^*-dependent inactivation of *Gcn5* in B cell lineages. The levels of CSR for WT, *Gcn5^f/f^, Gcn5^f/f^Pcaf^-/-^* are not significantly different (n.s.); the levels for *Gcn5^-/-^* and *Gcn5^-/-^Pcaf^-/-^* are lower than the former three groups (p=0.0030 and p=0.0121, correspondently, when compared to WT2); *Gcn5^-/-^* vs *Gcn5^-/-^Pcaf^-/-^* levels are similar (p=0.9997), and the *Aid^-/-^* has only background levels, WT2 vs *Aid^-/-^* is (p>0.0001).

## 4. Discussion

Both GCN5 and PCAF are involved in chromatin modification and DDR response, which made them relevant candidates to facilitate lymphocyte development (13, 14). One challenge was the lack of a relevant *in vivo* model because GCN5 and PCAF have certain redundant functions in acetylating H3K9, and because the germline inactivation of *Gcn5* results in early embryonic lethality in mice (32). Here, we generated and analyzed a complex mouse model which allowed studying of double-deficient *Gcn5^-/-^Pcaf^-/-^* B cells development *in vivo* and *ex vivo*. We used a germline knockout of *Pcaf* (32), a conditional knockout of *Gcn5^f/f^*, a knockin of CRE recombinase expressed under the B cell-specific *Cd19* promoter (34), and a knockin of YFP to track the activity of CRE recombinase (35).

For such a complex mouse model (*Gcn5^f/f^Pcaf^-/-^Cd19^+/cre^Rosa-26-YFP^+^*), multiple controls were used. In one line of the controls, the mice lacking PCAF, having floxed *Gcn5* gene but expressing no CRE recombinase were considered (*Gcn5^f/f^Pcaf^-/-^Rosa-26-YFP^+^*, Figure S2). The GCN5-deficient and GCN5/PCAF double-deficient B cells possessed developmental delay with lower levels of mature B cells in spleen and blood, and accumulation of progenitor B cells in bone marrow (Figures 1-4). Contrary, the control mice without CRE expression demonstrated WT levels of B cell development in all the groups, i.e. WT levels of B220+IgM+ mature B cells in the spleen (Figure S2 A and B) and blood (Figure 2S C and D). In addition, these mice possessed high levels of B220+IgM+ cells (Figure S2 E, F), as well as stable and low levels of pro-B cells (B220+IgM-CD43+) in bone marrow (Figure S2, G, and H).

The mice lacking PCAF and with conditional knockout of *Gcn5* in B cells were alive and resembled WT littermates (Figure 1A). One clear feature the *Gcn5^f/f^Pcaf^-/-^Cd19^+/cre^Rosa-26-YFP^+^* mice had was a small spleen (Figure 1 B and C), which was also the case in mice lacking only *Gcn5* in B cells. The small spleen could indicate a defect in B cell development, and we indeed found low numbers of mature B cells in the spleen, blood, and bone marrow. One could propose that mature B cells lacking GCN5 or both GCN5 and PCAF possess low proliferation speed or tend to trigger apoptosis. Alternatively, GCN5 and PCAF might be required for the V(D)J recombination. This option could be tested by, for example, using vAbl pre-B cell lines as we and others did before, e.g. (4, 15, 18, 23, 27–31). Another intriguing question is whether the physical presence or enzymatic activity of GCN5 and PCAF are required for the observed phenotypes, i.e. abrogated B cell maturation and reduced levels of CSR. To investigate this question one could use specific inhibitors of GCN5 and PCAF enzymes, or enzyme-dead mutations introduced to the *Gcn5* and *Pcaf* genes.

Inactivation of *Gcn5* in murine B cells also resulted in reduced lymphomagenesis in mice overexpressing MYC oncoprotein (44). Our findings further highlight this observation, suggesting that GCN5, and potentially also PCAF, enzymes are attractive targets for cancer therapy (44).

The CSR levels were reduced in B cells lacking GCN5 (Figure 4). The challenge in this set of experiments was that the mice of *Gcn5^f/f^Pcaf^-/-^Cd19^+/cre^Rosa-26-YFP^+^* genotype were rather rare and possessed a very low number of suitable B splenocytes (Figures 1 and 2). Although our data on IgG3 is sufficient, one could extend the study in the future by generating knockout cell lines lacking GCN5 and PCAF and suitable for CSR. One possible model system is CH12F3 cells capable to support CSR to IgA (45), which were used in the past for this kind of experiment (24, 37, 42). The CSR itself is a complex multistage process. Generating relevant cell lines will also provide tools to determine specific stages of CSR affected in GCN5-deficient mice, i.e. germline transcription, AID recruitment, generation of DSBs, or DNA repair.

## 5. Conclusions

Acetyltransferases GCN5 and PCAF possess redundant functions in B cell maturation. GCN5 is required for robust class switch recombination *ex vivo*.

## Author Contributions

Conceptualization, methodology, validation, formal analysis, investigation, resources, data curation – J.A.D., D.S., and V.O.; writing—original draft preparation, V.O.; writing—review and editing, V.O. and J.A.D.; supervision, J.A.D.; project administration, J.A.D. and V.O.; funding acquisition, J.A.D. and V.O. All authors have read and agreed to the published version of the manuscript.

## Acknowledgments

*Gcn5*-flox and *Pcaf*-null mice were kindly provided by the Sharon Dent laboratory.

## Institutional Review Board Statement

All experiments were performed in compliance with the Danish Working Environment Authority, the Danish Animal Experiment Inspectorate, the Department of Experimental Medicine (University of Copenhagen), and the Animal Resources Care Facility of Norwegian University of Science and Technology (NTNU, Trondheim, Norway), FOTS #11931 (2017) and FOTS #13405 (2018).

---

**Supplementary Figure S1.**
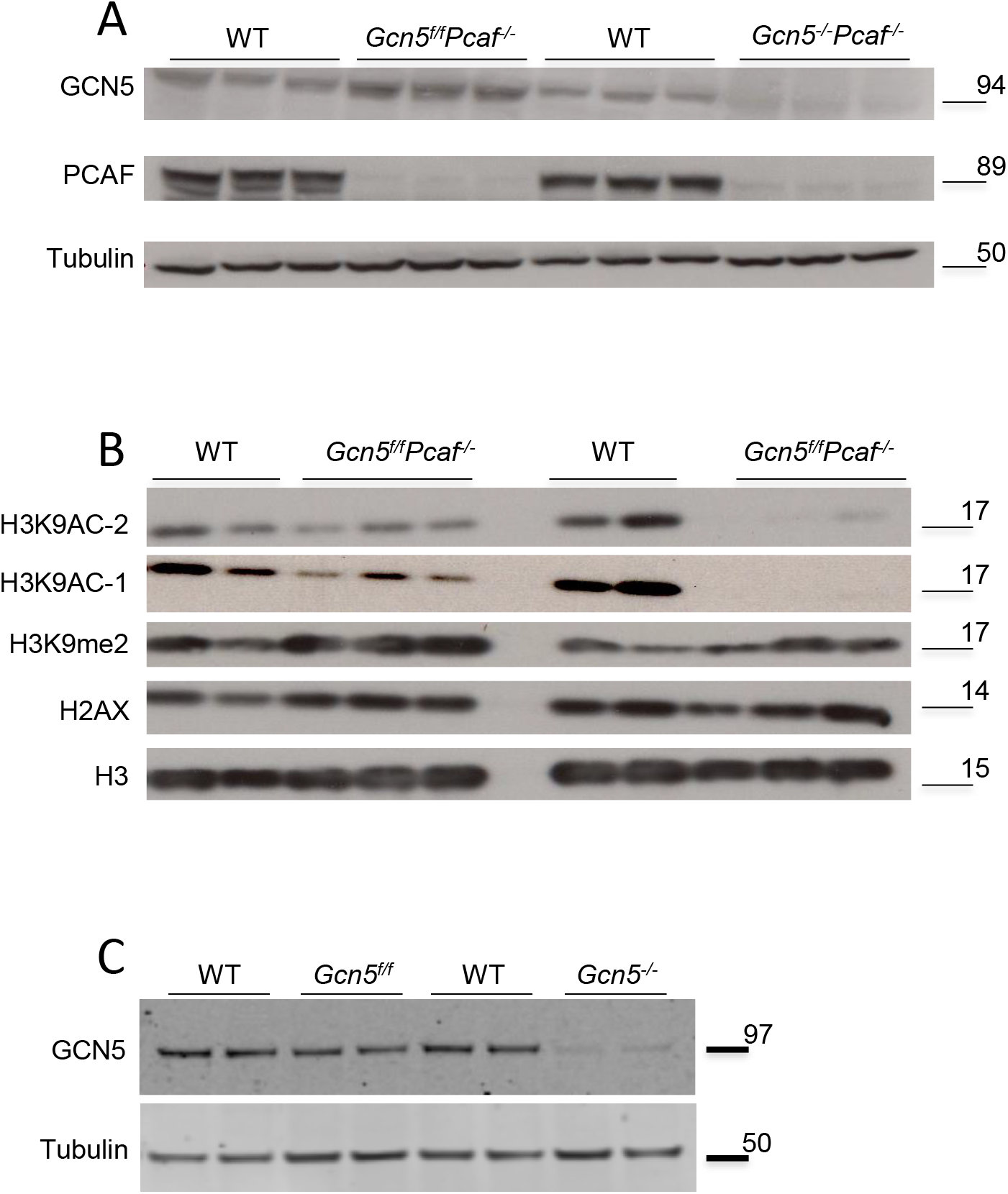

**Supplementary Figure S2.**
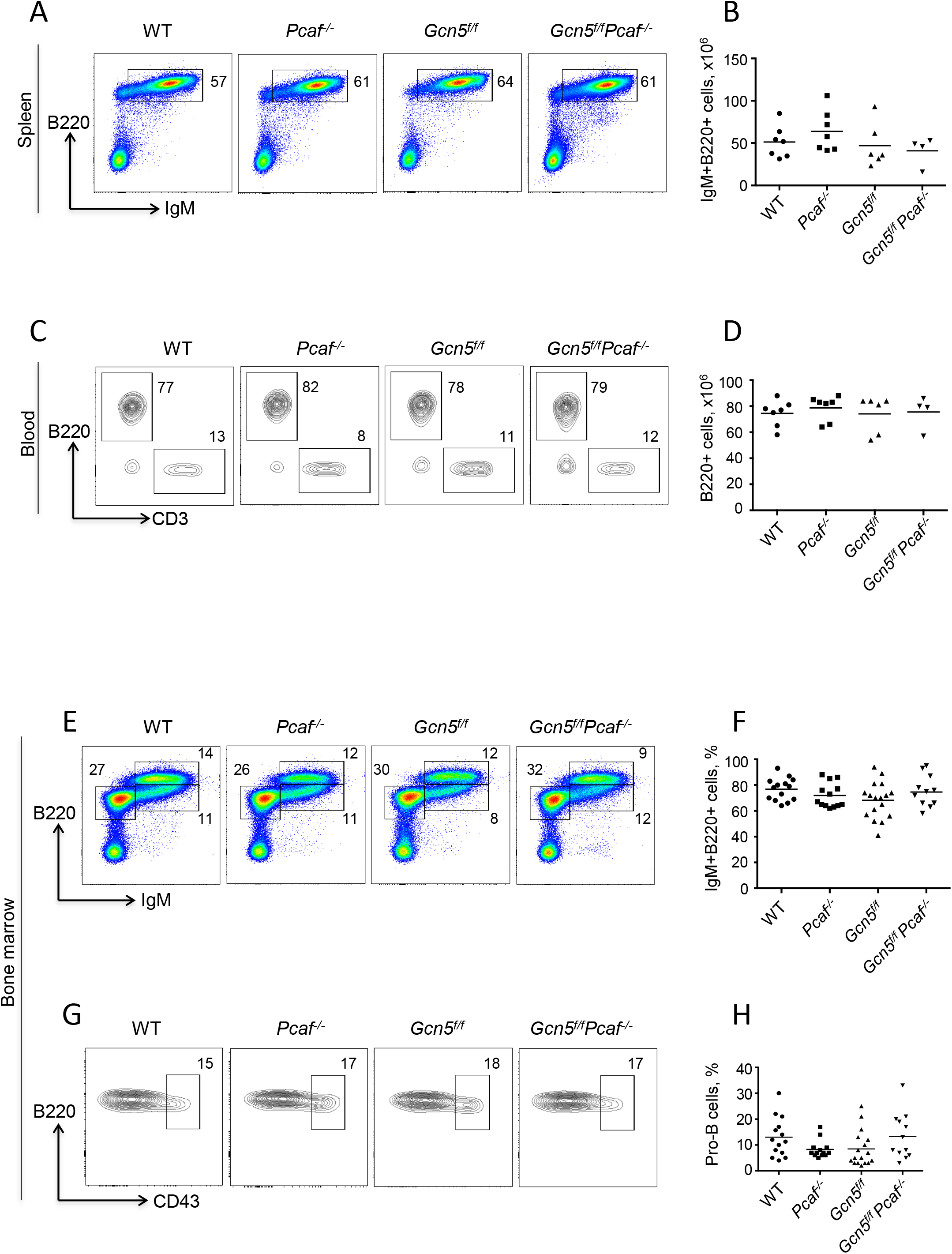

**Supplementary Figure S3.**
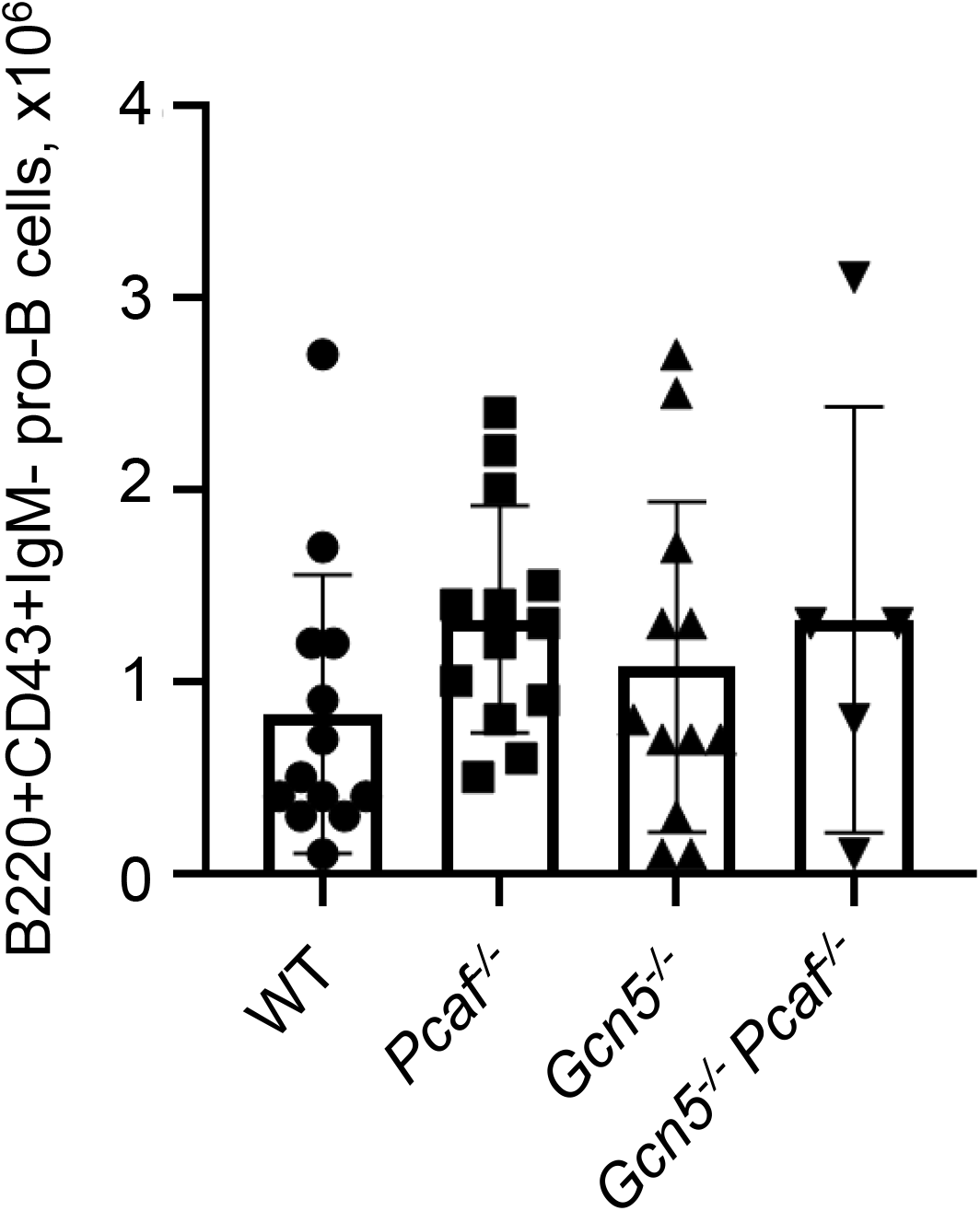

