## Supplementary material for "Acetyltransferases GCN5 and PCAF are required for B lymphocyte maturation in mice": Figures S1, S2, S3

Supplementary Figures 1 and 2

Aceltransferases GCN5 and PCAF are required for B cell maturation in mice

Valentyn Oksenych^1,2,3,4,5,^*

**Citation:**

Academic Editor: Firstname Lastname

Received: date

Accepted: date

Published: date

**Publisher’s Note:** MDPI stays neutral with regard to jurisdictional claims in published maps and institutional affiliations.

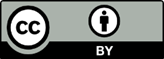

**Copyright:** © 2021 by the authors. Submitted for possible open access publication under the terms and conditions of the Creative Commons Attribution (CC BY) license (https://creativecommons.org/licenses/by/4.0/).

^1^	Department for Cancer Research and Molecular Medicine (IKOM), Norwegian University of Science and Technology, Laboratory Center, Erling Skjalgssons gate 1, 7491 Trondheim, Norway;

^2^	KG Jebsen Centre for B Cell Malignancies, Institute of Clinical Medicine, University of Oslo, N-0316 Oslo, Norway;

^3^	Institute of Clinical Medicine, University of Oslo, 0318 Oslo, Norway;

^4^	The NNF Center for Protein Research, Faculty of Health and Medical Sciences, University of Copenhagen, Blegdamsvej 3B, 2200 Copenhagen, Denmark

^5^	Department of Biosciences and Nutrition (BioNut), Karolinska Institutet, 14183 Huddinge, Sweden

**Supplementary Figure S1. Detection of GCN5, PCAF and histones using western blot**

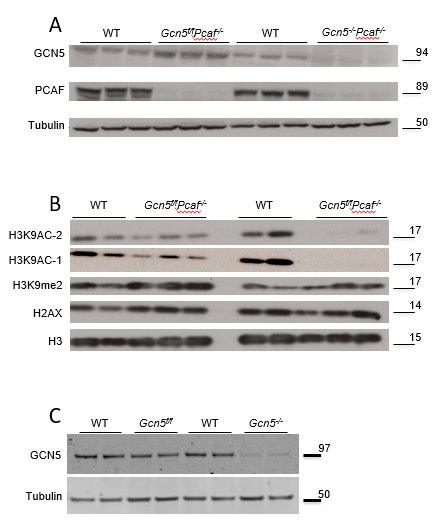

**Supplementary Figure S1. Detection of GCN5, PCAF and histones.**

**(A)** Detection of GCN5 and PCAF in primary murine B cells of indicated genotype. GCN5 signal is absent in the *Gcn5^-/-^* cells. PCAF signal is absent in the *Pcaf^-/-^* cells. Tubulin was used as a loading control.

**(B)** Western blot detecting Histones H3, H2AX, H3K9me2, H3K9ac (two different antibodies). H3K9Ac signal is missing in the cells lacking both GCN5 and PCAF.

**(C)** Detection of GCN5 and PCAF in primary murine B cells of indicated genotype. GCN5 signal is absent in the *Gcn5^-/-^* cells.

**Supplementary Figure S2. Detection of developing B cells in mice of indicated genotypes using flow cytometry**

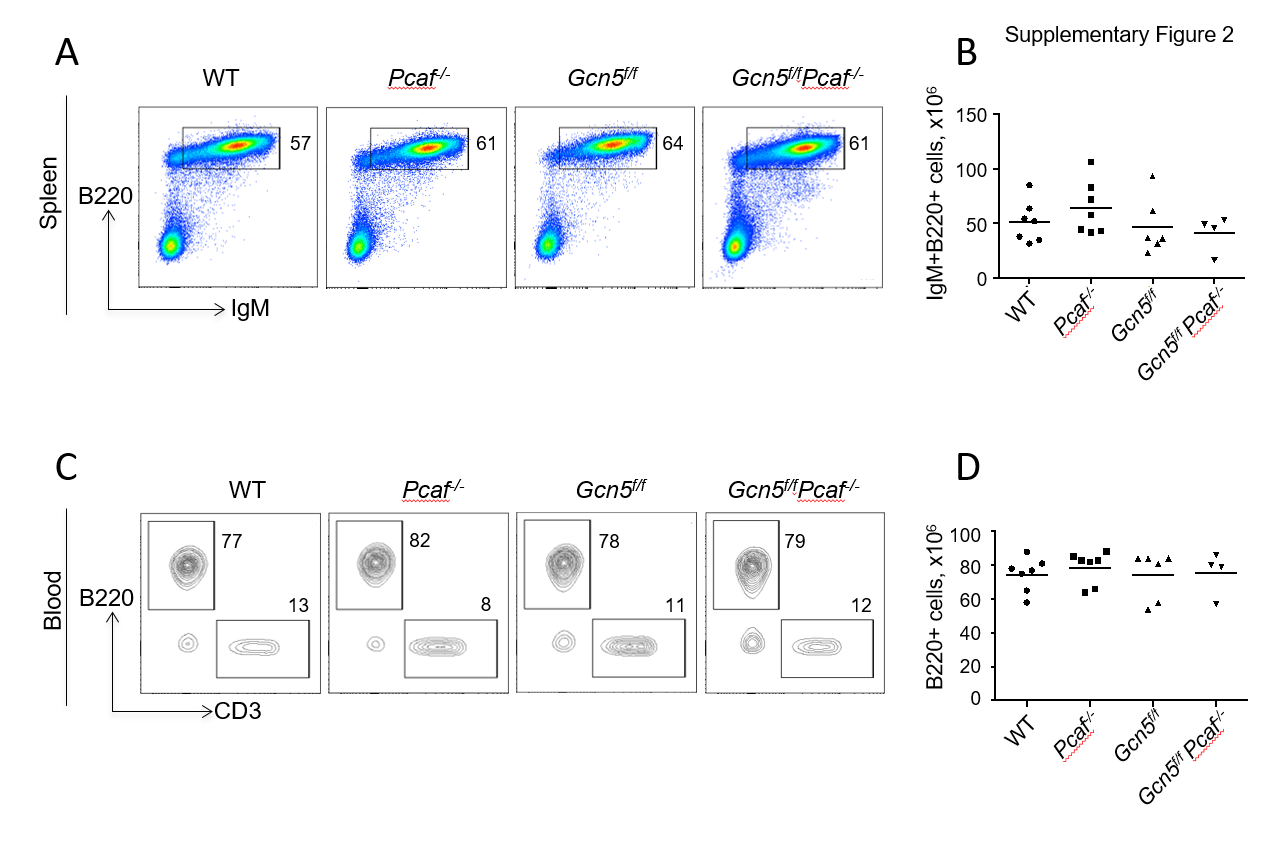

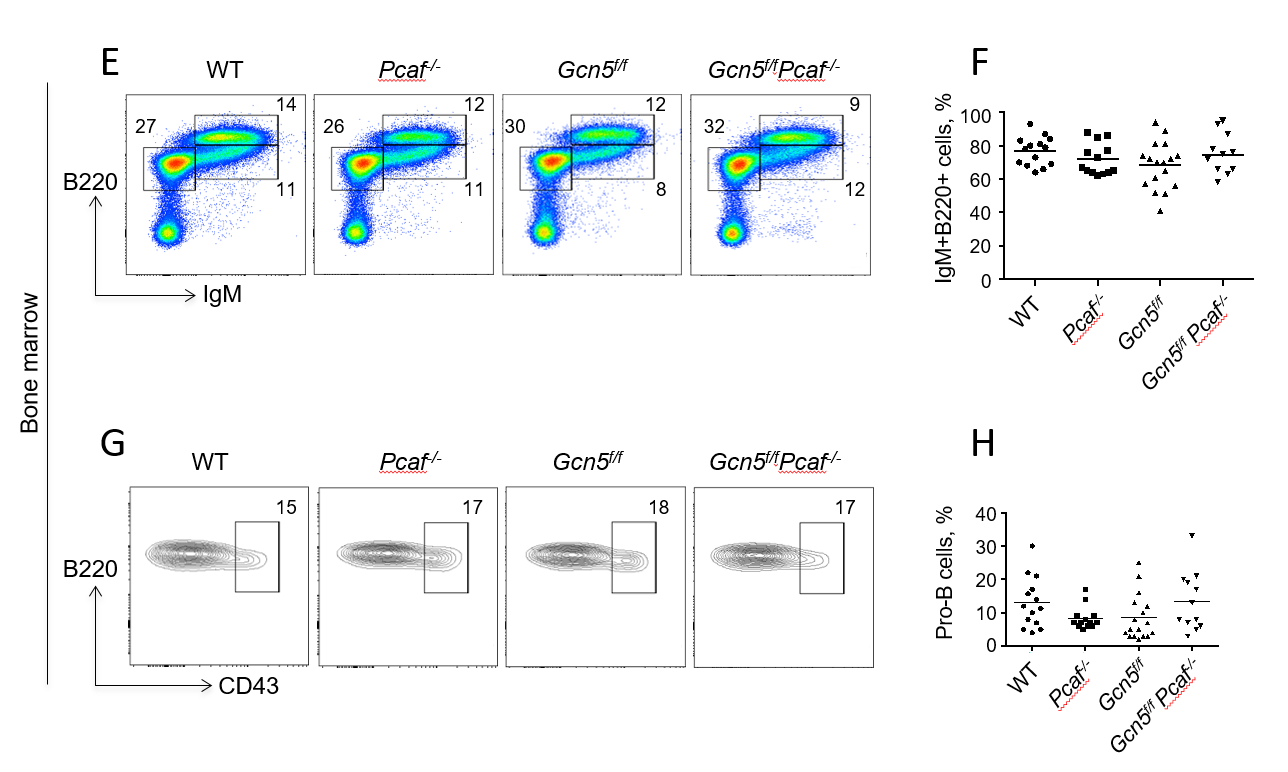

**Supplementary Figure S2. Detection of developing B cells in mice of indicated genotypes using flow cytometry**

(A) Example of flow cytometry detecting B220+IgM+ mature B cells in spleen.

(B) Summary of several experiments shown in (A).

(C) Example of flow cytometry detecting B220+ B cells and CD3+ T cells in blood.

(D) Summary of several experiments shown in (C).

(E) Example of flow cytometry detecting B220+IgM- and B200+IgM+ developing B cells in bone marrow.

(F) Summary of several experiments shown in (E).

(G) Example of flow cytometry detecting B220+IgM-CD43+ pro-B cells and B220+IgM-CD43- pre-B cells in bone marrow.

(H) Summary of several experiments shown in (G).

**Supplementary Figure S3.** Count of WT, *Pcaf*-deficient, *Gcn5*-deficient, and *Gcn5/Pcaf* double-deficient pro-B cells in bone marrow (million cells); n.s. p>0.3830

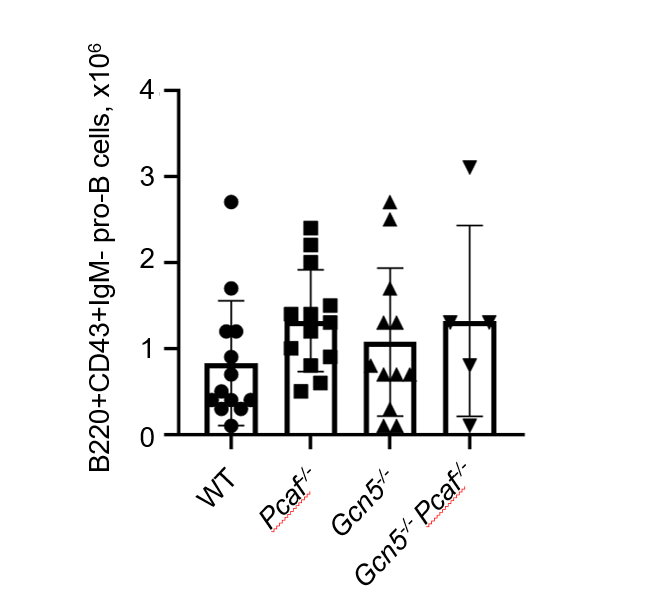
